# Automated high-throughput patch clamp electrophysiology of hiPSC-derived neuronal models

**DOI:** 10.1101/2025.05.12.653142

**Authors:** Federica Farinelli, Isaac Ostlund, Srinidhi Rao Sripathy, Debamitra Das, Gina Shim, Sangho Myung, Richard E. Straub, Brady J. Maher

## Abstract

The advent of human induced pluripotent stem cells (hiPSCs) and their differentiation into neurons and brain organoids has revolutionized our ability to model brain disorders in a human context. However, current technologies to assay the electrophysiological properties of human neurons in these models remain limited by throughput, as single-cell manual patch clamp is laborious and resource intensive. Here, we provide methods to perform high-throughput automated patch-clamp (APC) on hiPSC-derived neurons. We describe how to dissociate and perform voltage-clamp recordings on human neurons from three well-established protocols - 2D directed differentiation of cortical neurons, NGN2-induced neurons, and 3D cortical organoids - using the Nanion Syncropatch 384, a commercially available high-throughput APC system. Using this approach, we investigated the biophysical properties of voltage-gated sodium channels (VGSCs) and provide direct comparisons between manual and APC recordings across all three hiPSC-derived model systems. We demonstrate the capability of this automated system for pharmacological analysis of native human VGSC isoforms, which will enable compound screening approaches. Lastly, we provide methods to sort specific cellular populations within these hiPSC models using fluorescence-activated cell sorting (FACS) followed by APC. These methods and results provide a transformative and novel high-throughput technique for quantifying passive and active membrane properties in cell-type specific and/or genetically modified hiPSC-derived neurons.

## INTRODUCTION

The human-specific nature of nervous system disorders is a major barrier towards improving our understanding of pathophysiology, disease mechanisms, and the development of effective therapies. Often these disorders are highly complex, polygenic, and heterogeneous, with cognitive and clinical features that can only be effectively studied in humans. The recent advances in stem cell biology and the advent of human induced pluripotent stem cells (hiPSCs) has opened the door to modeling the human nervous system directly from patients ^1^. This exciting new paradigm has immense promise to improve our understanding of disease etiology and mechanisms, which should benefit their translation into effective therapies in the clinic.

The ability to derive and study specific cell types and tissues of the nervous system from hiPSCs is rapidly improving. A variety of differentiation protocols now exist to generate cell types that are specific to different regions of the central and peripheral nervous systems. Differentiation protocols are also capable of producing three-dimensional (3D) tissue (organoids), which can be used to study almost any nervous system structure ^2^. Through these advances in hiPSC differentiation, we can now generate models of almost any disorder in a patient-specific manner, which should improve development of not just disorder-specific but potentially patient-specific therapies (i.e. personalized medicine) ^3^.

To accomplish these lofty but critical goals, it is imperative to continue to improve upon current human models that more faithfully approximate the development, structure, and function of the cells and tissues they are meant to represent. To accelerate discovery and to untangle the polygenicity and heterogeneity inherent to most human-specific disorders, further technological advancement is needed to both generate and assay these human models with high-throughput.

In this paper, we describe a protocol for high-throughput APC electrophysiology on human neurons derived from hiPSCs. Patch-clamp electrophysiology is the gold standard method for quantifying the physiological properties of excitable cells ^4^. However, its widespread use in hiPSC-derived models has been limited due to its highly technical nature and low throughput. We have made substantial progress on these limitations by optimizing the use of hiPSC-derived neurons with a commercially available APC platform. We provide detailed optimized protocols to record from three different commonly used differentiation protocols. These include co-cultures of 2D neurons on rodent astrocytes, NGN2-induced neurons, and 3D cortical organoids (hCOs). We studied intrinsic membrane properties and voltage-gated sodium channel (VGSC) currents across these hiPSC-derived neuronal models and compared the results from automated and manual patch-clamp. We demonstrated that the isoform-selective VGSC inhibitor XPC6444 partially blocks native VGSC currents in human neurons. Lastly, we demonstrate that the addition of a fluorescence-activated cell sort (FACS) step to the protocol increases recording efficiency and allows for APC recording of specific cell types and/or genetically manipulated neurons.

## METHODS

### hiPSC cell lines

Generation and validation of the 16 hiPSC lines used in this study are described in detail in Page et al. 2022^5^. All hiPSCs were grown as feeder-free cultures fed with Stemflex media for maintenance.

### Differentiation into 2D cortical neurons

hiPSCs were differentiated into 2D cortical neurons as previously described ^6^ with modifications. The modifications carried out were as described in Page et al. 2022 ^5^. 2D cortical neuron cultures were cocultured with rat astrocytes and grown on poly-d-lysine/laminin-coated coverslips for 10 weeks (DIV 70). Neuronal cultures at 10 weeks were either used directly for manual patch-clamp or harvested & dissociated (as described below) for APC experiments.

### Differentiation into 3D cortical organoids (hCOs)

hiPSCs were differentiated into hCOs as previously described ^7^. Briefly, hiPSCs were dissociated into single cells using accutase and plated in AggreWell^TM^ 800 at 3×10^6^ cell density per well in Stemflex media supplemented with Rocki (Y-27623). After 48h, hCOs were carefully harvested using a 5ml serological pipette, and wells were flushed two additional times with Stemflex using a p1000 pipette to ensure the collection of all spheroids. Harvested spheroids were transferred to ultra-low attachment dishes in neural induction media (NIM; Essential 6 medium supplemented with dual-SMAD pathway inhibitors - Dorsomorphin (2.5uM), SB-431542 (10uM) and XAV (2.5uM)). Media changes were performed daily from Day 2 to Day 5 with NIM media. On the 6th day in suspension, the media was changed to neural differentiation media (NDM); a combination of Neurobasal-A media, B27 supplement without vitamin A, Glutamax, and Penicillin/Streptomycin). The neural media was supplemented with EGF (20ng/ml) and FGF (20ng/ml) for 19 days, with daily media changes for the first 10 days, and every other day from the 9 days after (until day 24). Following this, from Day 26 to Day 42, neural media was supplemented with BDNF (20ng/ml) and NT3 (20ng/ml) with media changes every other day. Following Day 43, media changes were done every 4 days with only neural differentiation media without any growth factors.

### Differentiation of NGN2-induced neurons

hiPSCs were transduced with commercially available Neurogenin2 (Ngn2) and reverse tetracycline-controlled trans-activator (rtTA) lentivirus using polybrene as previously described ^8,9^. Infected cells were subsequently treated with doxycycline to induce expression of Ngn2 and rtTA virus and selected using puromycin to collect only NGN2-expressing cells. Surviving cells were replated on matrigel-coated 6-well plates at 1×10^6^ cells/well density and on 24-well plates with coverslips at 100K cells/well density and differentiated for 21 days (DIV 21). Neuronal cultures at day 21 were either directly used for manual patch-clamp or harvested & dissociated for APC experiments.

### Single-cell dissociation of hiPSC-derived neurons for Nanion multipatch

To prepare cells for Nanion automated patching, a modified version of the Worthington Papain Dissociation kit (Worthington, Cat # LK003150) manufacturer’s protocol was used ^10^. As per the protocol, before applying Papain, DNAse and Ovomucoid inhibitor were reconstituted with Earle’s medium. Reconstituted DNase was mixed with reconstituted Papain and 10uM Rocki (Y-27623) was added to the solution to support survival of dissociated cells.

For hCO dissociation (Day 150), 2-3 hCOs per line were transferred to 60mm dishes. After removing any residual media, 2.5 ml of Papain/DNase/Rocki solution was added, and the hCOs were chopped into small pieces (<1mm) and incubated at 37°C for 30 mins with continuous shaking. After 30 mins, the organoid pieces were further dissociated by trituration with a p1000 pipette (pipette 3-4 times) and returned to incubate at 37°C for an additional 10 mins.

For 2D cortical neurons (DIV 70) and NGN2-induced neurons (DIV 21), either 1ml/well for a well of 6-well plate (NGN2-neurons) or 500ul/well for a well of 24-well plate (2D cortical neurons) of the Papain/DNAase/Rocki solution was added and incubated for an hour at 37°C. After one hour, 1/10^th^ volume (100ul or 50ul) of inhibitor solution was added to each well and triturated gently using a P1000 pipette to break the neurons into single cells in suspension.

The dissociated cell suspension was gently overlaid dropwise onto a 4X volume of inhibitor solution, where it separated into two distinct layers of inhibitor solution and dissociated cell suspension. To prevent damaging cell membranes, care was taken to gently triturate/pipette cells during dissociation, and they were frequently monitored under a microscope. Dissociated cells were then centrifuged at 200g for 5 minutes at gradual acceleration and deceleration (6 & 6). The supernatant was removed, and the cell pellet was resuspended in 1 ml of Nanion extracellular recording solution. Cells were counted, and a cell suspension (100K/mL) was transferred to the cell reservoir of the APC system.

### Automated patch-clamp (APC)

All APC recordings were performed using the SyncroPatch 384i (Nanion Technologies). Freshly dissociated cells were placed into the cell reservoir (∼10°C, 200 RPM shaker). Cells were then delivered to the recording chip via a liquid handling pipette. A catch pressure of −75 mBar was applied for 5s and then a holding pressure of −30 mBar was applied until the number of wells reporting a seal resistance greater than 10 MΩ reached equilibrium (generally two minutes, with 60-85% wells reaching 10 MΩ seal resistance). To improve the seal, cells were exposed to a divalent-enriched external solution (seal enhancer) containing 5 mM CaCl₂ until the captured cells achieved a seal resistance of 300 MΩ, followed by two washes with a standard external solution (described below). The cells were incubated for 3 minutes in a standard external solution while a cesium fluoride internal solution containing 8 µM escin was applied, followed by an additional 3-minute incubation to establish whole-cell configuration before executing voltage-clamp protocols. All electrophysiological protocols and data were created and digitized using PatchControl 384 software (Nanion Technologies). Recordings were performed on single-hole S-type chips (Nanion Technologies). Each well of the SyncroPatch 384 has an individual headstage and amplifier, and can record unique cell traces, therefore, each well is denoted as *n* = 1. Biophysical parameters such as seal resistance, capacitance, and series resistance were determined from each well after the application of a test pulse. All parameters are monitored over time and can be recorded for individual experiments. VGSC currents were evoked by a voltage-step protocol from a holding potential (*V*_hold_) of −100 mV. The protocol produced 50ms steps ranging from −80mV to 30 mV, followed by a return to *V*_hold_. Peak current amplitude was recorded and normalized using cell capacitance. Steady-state inactivation was induced using a series of 400 ms conditioning steps ranging from −130 mV to −20 mV, followed by a 20ms test step to −20 mV. The inactivated currents were normalized to the maximal current using the DataControl 384 software.

### APC and manual patch-clamp recording solutions

The intracellular recording solution for APC contained (in mM) 110 CsF, 10 CsCl, 10 NaCl, 10 EGTA, 10 HEPES (pH 7.3). The standard extracellular recording solution for APC contained (in mM) 140 NaCl, 4 KCl, 2 CaCl_2_, 1 MgCl2, 5 Glucose, 10 HEPES (pH 7.3). The seal enhancer solution contained (in mM) 140 NaCl, 4 KCl, 5 CaCl_2_, 0.5 ml MgCl_2_, 5 Glucose, 10 HEPES (pH7.3). The intracellular solution for manual patch-clamp of 2D, 3D, and NGN2 neurons (in mM) 10 NaCl, 140 CsF, 10 HEPES, 1 EGTA, 15 D-glucose, and TEA-Cl (pH 7.3). The extracellular recording buffer for manual patch-clamp of 2D and NGN2 neurons (in mM) 40 NaCl, 75 choline-Cl, 1 CaCl_2_, 1 MgCl_2_, 10 HEPES, 20 TEA-Cl, 1 4AP, 0.1 CdCl and 10 D-glucose (pH 7.3). The extracellular solution for manual patch-clamp of 3D neurons (in mM) 125 NaCl, 1 CaCl_2_, 1 MgCl_2_, 10 HEPES, 20 TEA-Cl,1 4AP, 0.1 CdCl and 10 D-glucose (pH 7.3).

### Intracellular perfusion

The intracellular perfusion system is divided into 12 channels, each serving 32 wells. Intracellular solution is supplied by 1 L glass bottles by applying negative pressure underneath the fluidic system. The minimum volume for filling one channel is roughly 4 ml (∼50 ml for each chip run, including wash cycles).

### Pharmacology

XPC-6444 (Medchemexpress LLC) is a highly potent, isoform-selective, and CNS-penetrant NaV1.6 (IC50 = 41 nM for hNaV1.6) and NaV1.2 inhibitor (IC50 = 125 nM). To ensure complete inhibition of both human isoforms, we used a final concentration of 250 nM. To minimize membrane property alterations due to liquid exchanges, XPC-6444 was prepared at 5X the desired concentration, and 10uL was added to each well containing 40uL of external standard. VGSC currents were fully blocked by 1 μM TTX.

### Quality control parameters

Wells must pass several quality control (QC) parameters, which include board check, contact, and liquid junction potential, and cells must pass several QC parameters such as *R*_Seal_ > 300 MΩ, Capacitance > 1pF, and *I*_peak_ > 100 pA. These filters were applied automatically after recording in the DataControl 384 software. Cells that failed to meet these cutoffs were removed from further analysis. To avoid dynamic and steady-state errors, we only accepted cells with series resistance <40 MΩ.

### Data analysis

The SyncroPatch 384 platform has a software package consisting of PatchControl 384 (for data acquisition) and DataControl 384 (for data analysis; both Nanion Technologies). Raw data were further filtered based on the quality of their Boltzman fit and the corresponding values were exported using the “Export to Excel” function of DataControl 384 and wells that failed automated QC filters were removed. Results were then compiled and plotted using GraphPad 10.

## RESULTS

The basic workflow for obtaining APC recordings from hiPSC-derived neurons is outlined in **Figure 1**. We differentiated hiPSCs into neurons or organoids using previously published protocols ^5,7,8^. Cultures were grown for appropriate amounts of time to allow for neurons to mature. Cells were dissociated using a commercially available papain dissociation kit. Dissociated cells were diluted to a final cell volume of 100K/ml (range 15K/mL to 100K/mL; see below for FACS) in extracellular recording buffer and transferred to the APC systems. To improve recording success, care must be taken during dissociation to keep cell membranes healthy and to clear cellular debris. Cells were dispensed across the 384 well plate using the robotic pipette liquid handler. Each well contains a single pore for capturing/catching and break-in of a single cell, which requires customized suction settings for each type of dissociated cell. Similar to manual patch-clamp, a high resistance seal is required before whole-cell break-in. A catch pressure of −75 mBar was applied for 5s, and then a holding pressure of −30 mBar was applied until the number of wells reporting a seal resistance greater than 10 MΩ reached equilibrium (generally two minutes, with 60-85% cells reaching 10 MΩ seal resistance). To improve the sealing of the cell to the pore, a high Ca^2+^ containing seal enhancer solution was applied to the extracellular side of the cells for 1 minute before washing out with standard extracellular buffer. After cell catch, we found it difficult to use a suction protocol to reliably break-in to cells while maintaining a high enough series resistance for adequate voltage-clamp. Instead, we had success with perforated patch by adding escin (8 µM), a mixture of saponins that produces pores in cell membranes ^11,12^, to the intracellular solution throughout the length of the recording. This approach greatly improved break-in, series resistance, and our success rate.

**Figure 1.**
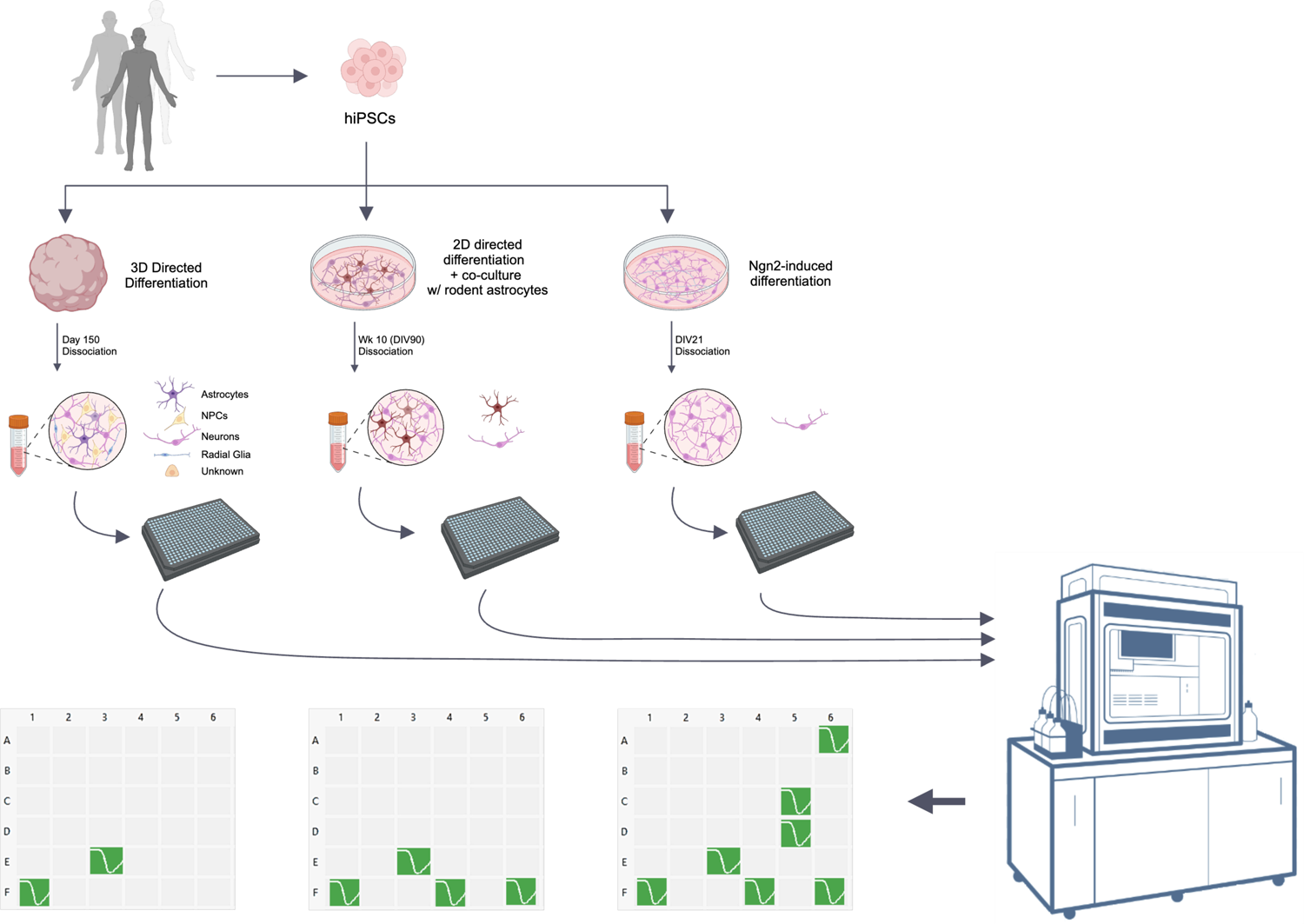
Workflow from hiPSC cells to high-throughput automated patch clamp recordings. Donor cells are reprogrammed into hiPSCs. hiPSCs are differentiated into cortical organoids, co-cultures of human cortical neurons and rodent astrocytes, or NGN2-induced neurons. hiPSC-derived neurons are then dissociated, counted, and dispensed into the automated patch clamp system for whole-cell electrophysiological recordings.

Following this whole-cell break-in procedure, neurons were voltage-clamped at −70mV, and a series of voltage-clamp protocols was applied to quantify the neuronal membrane properties and the activation and inactivation kinetics of VGSC currents (**Figure 2**). For steady-state VGSC activation, currents were evoked from a holding potential (Vhold) of −100 mV by a series of 5 mV (50 ms) voltage steps ranging between −80 to +30 mV followed by a return to Vhold (**Figure 2a**). We then applied a steady-state VGSC inactivation protocol that consisted of a series of conditioning voltage steps from −130 to 20 mV (400 ms) followed by a −20 mV (20 ms) test voltage step (**Figure 2b**). For every recording, the acquisition software also reported capacitance, membrane resistance, and series resistance (**Figure 2c**). Through several rounds of dissociation and recording, we optimized the protocol for whole-cell break-in which resulted in significant increases in the efficiency of obtaining a successful recording, which was defined by the percent of wells recording from a cell with a VGSC current greater than 100 pA. Across the three hiPSC-derived models (2D, 3D, NGN2), using the same whole-cell break-in procedure, we observed a difference in our recording success rate (**Figure 2c**). The success rate appeared to depend on the cell-type complexity of the cultures that were being recorded. Organoids showed the lowest success rate (∼5%) which we primarily attributed to the cell-type diversity of this model. Cortical organoids contain many different non-neuronal and neuronal cell types (e.g. NPCs, astrocytes, and neurons) ^7^, each of which contribute to the total number of cells dispensed onto the APC system. 2D co-cultures showed an intermediate success rate (∼10%), being composed of approximately 20% neurons and 80% rat astrocytes ^5^. We observed the highest success rate (∼19%) from NGN2-induced neuronal cultures which generate a relatively pure population of human neurons ^8,13^.

**Figure 2.**
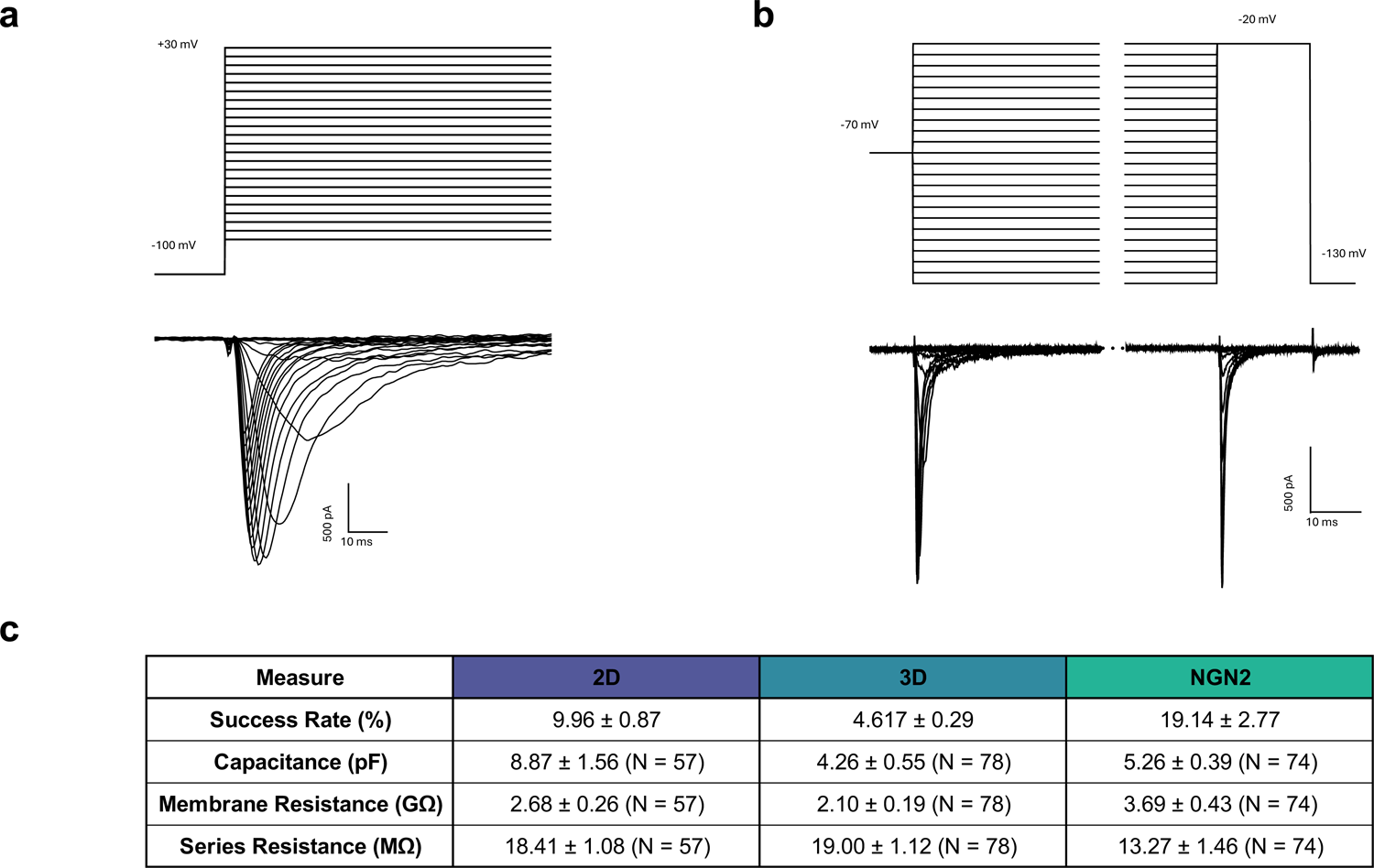
Example voltage-clamp recordings, success rate, and membrane properties of hiPSC-derived neurons recorded with the automated patch-clamp system. **(a)** VGSC activation protocol and representative electrophysiology traces from a 2D neuron. **(b)** VGSC inactivation protocol and representative electrophysiology traces from a 2D neuron. **(c)** Table of success rate and membrane properties obtained from three different hiPSC-derived neuronal models (2D, 3D, NGN2) using the automated patch-clamp system. Success rate is the percentage of wells with a successful recording. The N for capacitance, membrane resistance, and series resistance is the number of recorded neurons. All values are mean ± sem.

### Automated Electrophysiological Analysis of Membrane Properties and VGSC Currents

VGSCs are a primary source of excitability in neurons and are responsible for setting the membrane potential and for the generation of action potentials. Using manual patch-clamp, we previously identified differences in the intrinsic membrane properties and function of VGSCs in hiPSC-derived neurons from schizophrenia patients ^5^, which motivated the development of the high-throughput electrophysiology method described here. Using the APC system, we recorded whole-cell intrinsic membrane properties and VGSC currents in voltage-clamp mode from cortical neurons co-cultured with rat astrocytes in monolayer (2D; **Figure 3**), cortical organoids (3D; **Figure 4**), and NGN2-induced neurons in monolayer (NGN2; **Figure 5**).

**Figure 3.**
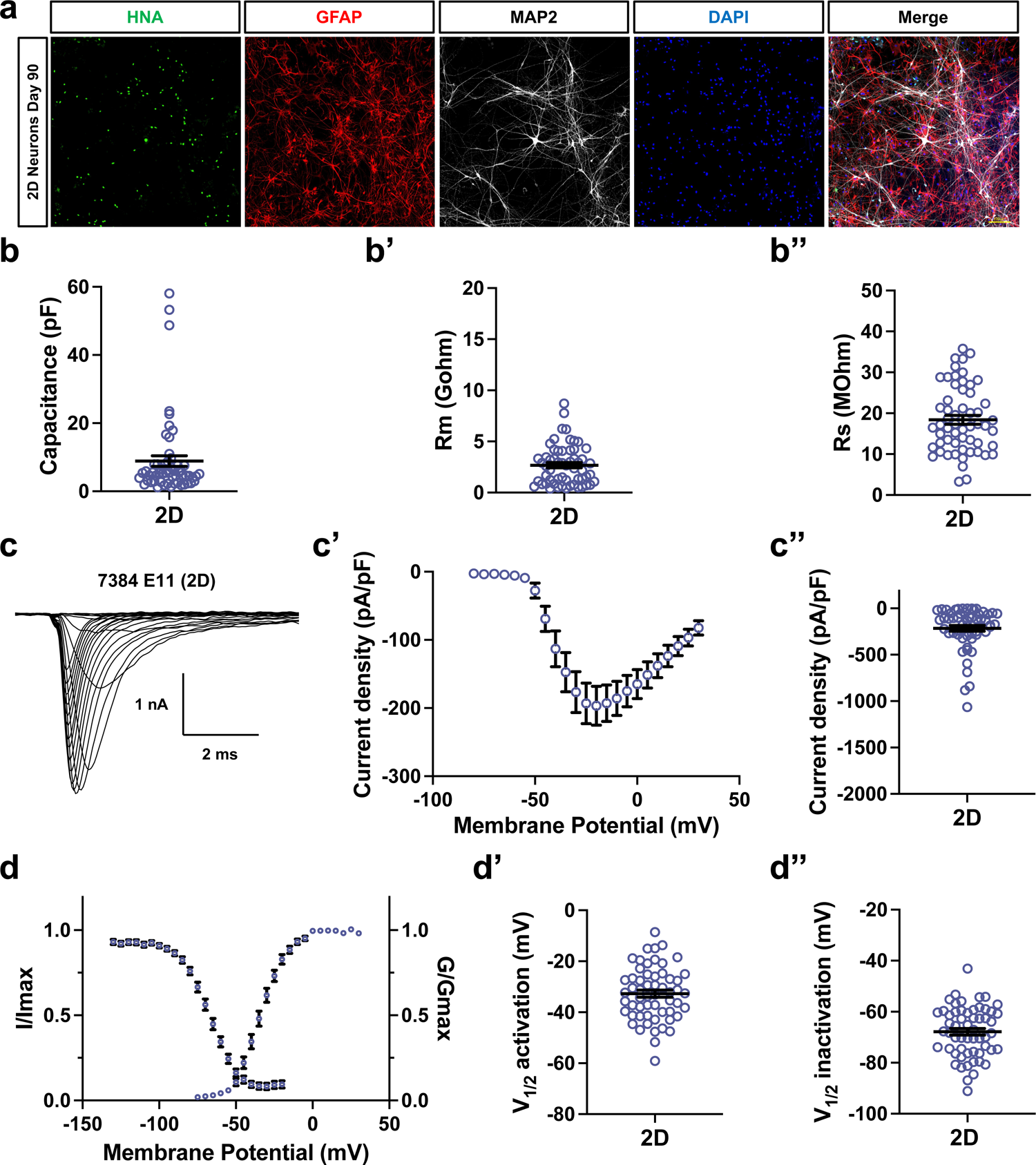
Automated patch-clamp of 2D human neurons co-cultured with rat astrocytes. **(a)** Representative images of 2D human neurons co-cultured with rat astrocytes. Day 90 cortical neurons are double-positive for HNA and MAP2. Rat astrocytes are positive for GFAP and negative for HNA. **(b)** Intrinsic membrane properties of neurons co-cultured in 2D monolayers with rat astrocytes. The average **(b)** capacitance is 8.87±1.56 pF (N=57), **(b’)** membrane resistance 2.68±0.26 GΩ (N=57), and **(b’’)** series resistance 18.41±1.1 MΩ (N=57). **(c)** Representative VGSC traces in response to voltage steps. **(c’)** Summary current voltage (I-V curve) plot **(c”)** Summary peak current density plot (−215.4±30.4 pA/pF (N=57)). **(d)** Summary plot of steady state voltage dependence of VGSC activation (G/Gmax) and inactivation (I/Imax). **(d’)** Summary plot of V_1/2_ activation of VGSCs (V_1/2_=-32.71±1.38 mV (N=57)) **(d’’)** Summary plot of V_1/2_ inactivation of VGSCs (V_1/2_=-67.84±1.30 mV (N=57)).

**Figure 4.**
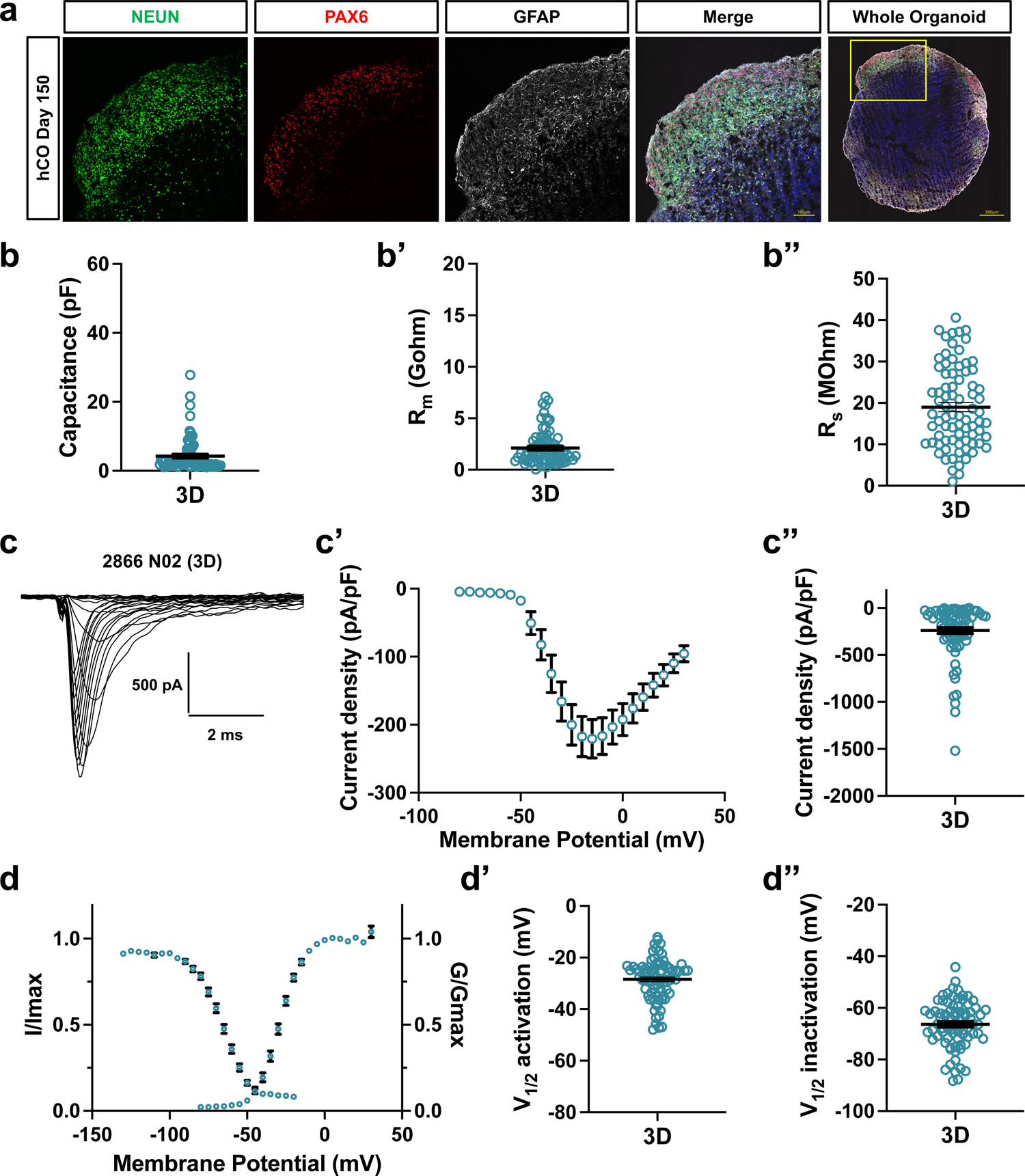
Automated patch-clamp of 3D cortical organoids. **(a)** Representative images of a day 150 cryosectioned cortical organoid. Cortical organoids contain NEUN-positive neurons, PAX6-positive NPCs, and GFAP-positive astrocytes. **(b)** Intrinsic membrane properties of neurons co-cultured in 2D monolayers with rat astrocytes. The average **(b)** capacitance is 4.26±0.55 pF (N=78), **(b’)** membrane resistance 2.10±0.19 GΩ (N=78), and **(b’’)** series resistance 19.00±1.12 MΩ (N=78). **(c)** Representative VGSC traces in response to voltage steps. **(c’)** Summary current voltage (I-V curve) plot **(c”)** Summary peak current density plot (−239.8±32.62 pA/pF (N=78)). **(d)** Summary plot of steady state voltage dependence of VGSC activation (G/Gmax) and in) activation (I/Imax). **(d’)** Summary plot of V_1/2_ activation of VGSCs (V_1/2_= −28.45±0.90mV (N=78)) **(d’’)** Summary plot of V_1/2_inactivation of VGSCs (V_1/2_= −66.36±1.06 mV (N=76)).

**Figure 5.**
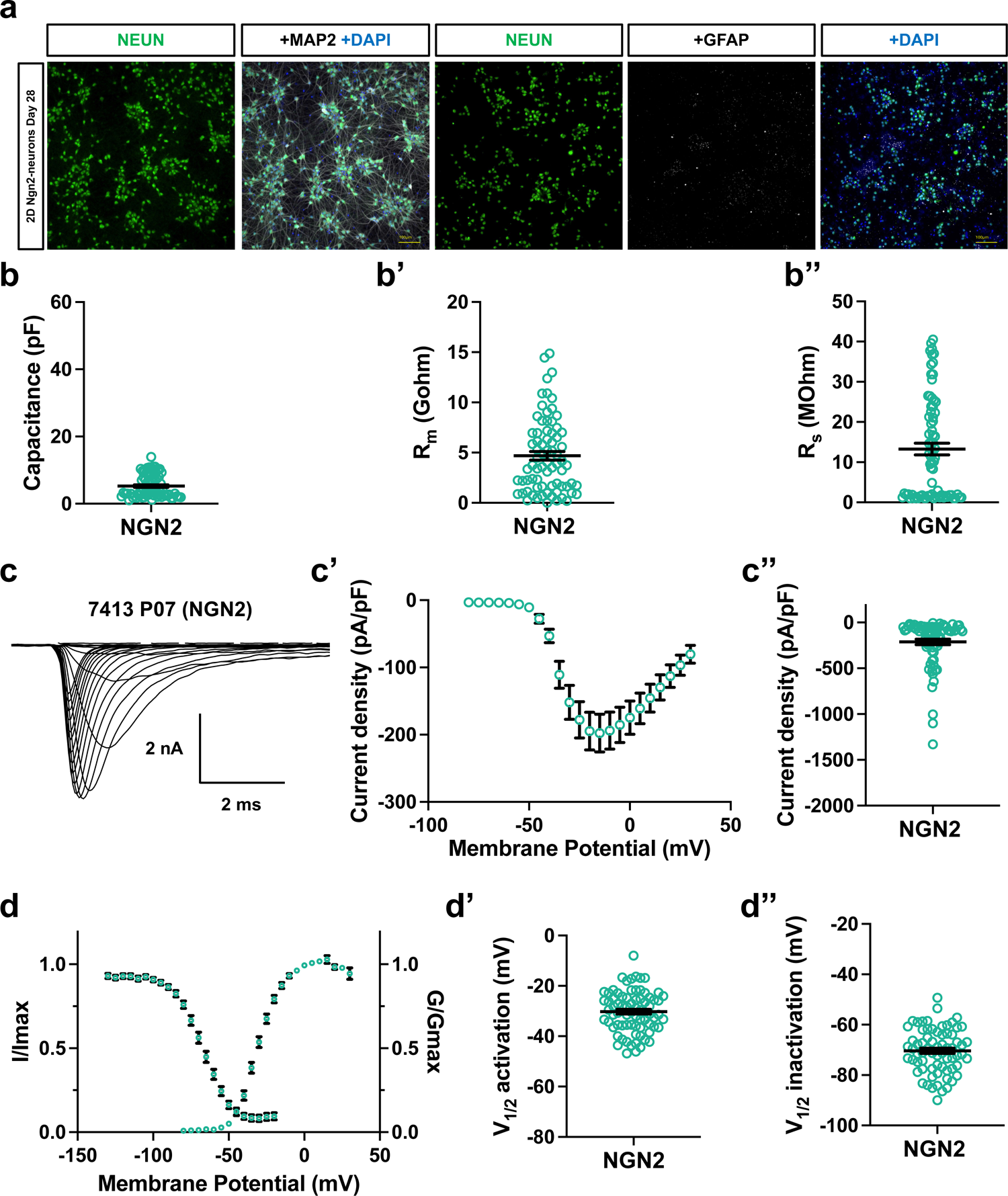
Automated patch-clamp of NGN2-induce neurons. **(a)** Representative images of day 28 NGN2-induced neurons. Neurons are co-labelled by NEUN and MAP2 and negative for the astrocyte marker GFAP. **(b)** Intrinsic membrane properties of neurons co-cultured in 2D monolayers with rat astrocytes. The average **(b)** capacitance is 5.26±0.39 pF (N=74), **(b’)** membrane resistance 4.69±0.43 GΩ (N=74), and **(b’’)** series resistance 13.27±1.46 MΩ (N=74). **(c)** Representative VGSC traces in response to voltage steps. **(c’)** Summary current voltage (I-V curve) plot **(c”)** Summary peak current density plot (−211.90 ±29.96 pA/pF (N=74)). **(d)** Summary plot of steady state voltage dependence of VGSC activation (G/Gmax) and in) activation (I/Imax). **(d’)** Summary plot of V_1/2_ activation of VGSCs (V_1/2_=-30.24±0.96mV (N=74)) **(d’’)** Summary plot of V_1/2_ inactivation of VGSCs (V_1/2_=-70.34±1.05 mV (N=69)).

### Automated recordings from dissociated cortical neurons co-cultured with rat astrocytes in monolayer (2D)

We generated co-cultures of cortical neurons and primary rat astrocytes using a previously established protocol ^5,6^. Immunostaining of these cultures demonstrates that they contain human cortical neurons co-labeled by MAP2 and human nuclear antigen (hNA), whereas rat astrocytes are positive for GFAP and negative for hNA (**Figure 3a**). We recorded from a single hiPSC cell line at day 70, across two dissociations. 2D neurons had an average membrane capacitance of 8.87±1.56 pF (n=57), membrane resistance of 2.68 ±0.26 GΩ (n=57), and series resistance of 18.41±1.08 MΩ (n=57) (**Figure 2c, 3b-b’’**). Neurons were then subjected to a voltage step protocol which led to the activation of VGSC currents (**Figure 3c**) that showed an average peak current density of −215.4±30.44 pA/pF (n=57) (**Figure 3c’, c”)**, and an average V_1/2_ activation of −32.71±1.38 mV (n=57) (**Figure 3d, d’**). A VGSC inactivation protocol was then applied and 2D neurons showed an average V_1/2_ inactivation of −67.84±1.30 mV (n=57) (**Figure 3d, d’’**).

### Automated recordings from dissociated cortical organoids (3D)

We generated cortical organoids using a previously established protocol ^7^. To characterize the cellular populations within these organoids, we immunostained cryosections and observed NeuN-positive neurons, GFAP-positive astrocytes, and PAX6-positive NPCs (**Figure 4a**). We recorded from 3D neurons across 12 different hiPSC lines. 3D neurons showed an average membrane capacitance of 4.26±0.55 pF (n=78), membrane resistance of 2.10±0.19 GΩ (n=78), and series resistance of 19.00 ± 1.12 MΩ (n=78) (**Figure 2c, 4b-b’’**). Neurons were then subjected to a VGSC activation protocol which led to the activation of VGSC currents (**Figure 4c**) that showed an average peak current density of −239.8 ±32.62 pA/pF (n=78) (**Figure 4c’, c”)**, and an average V_1/2_activation of −28.45±0.90 mV n=78 (**Figure 4d,d’**). A VGSC inactivation protocol was then applied and 3D neurons showed an average V_1/2_ inactivation of −66.36± 1.06 mV (n=76) (**Figure 4d, d’’**).

### Automated recordings from dissociated NGN2-induced neurons (NGN2)

We generated NGN2-induced neurons using a previously established protocol ^14^. Immunostaining of these cultures demonstrated that they contain human neurons that are co-labeled by MAP2 and NeuN, and these cultures were devoid of GFAP-positive astrocytes (**Figure 5a**). We recorded from NGN2 neurons derived from a single hiPSC line at day 21, across 3 dissociations. NGN2 neurons displayed an average membrane capacitance of 5.26±0.39 pF (n=74), membrane resistance of 4.69±0.43 GΩ (n=74), and series resistance of 13.27±1.46 MΩ (n=74) (**Figure 2c, 5b-b’’**). Neurons were then subjected to a VGSC activation protocol which lead to activation of VGSC currents (**Figure 5c**) that showed an average peak current density of −211.90±29.96 pA/pF (n=74) (**Figure 5c’, c”)**, and an average V_1/2_ activation of −30.24±0.96 mV (n=74) (**Figure 5d, d’**). A VGSC inactivation protocol was then applied and 2D neurons showed an average V_1/2_ inactivation of −70.34±1.05 mV (n=69) (**Figure 5d, d’’**).

### Comparison of intrinsic membrane and VGSC properties among hiPSC models

Intrinsic membrane and VGSC properties are developmentally regulated. As neurons mature, they typically increase in size and express more ion channels on their surface, which subsequently leads to an increase in capacitance, a decrease in membrane resistance, and larger VGSC currents ^15^. For hiPSC-derived neuronal models, differences in neuronal maturation are dependent on the differentiation protocol employed, the presence or absence of astrocytes, and the developmental time point studied (i.e., DIV). For the three models tested here, the maturation of 3D organoids most closely matches human brain development ^16^, whereas overexpression of NGN2 induces rapid neuronal maturation ^13^, and the addition of rat astrocytes to 2D neurons also accelerates their maturation ^17,18^.

When comparing intrinsic membrane properties, we observed that 2D neurons exhibited the largest capacitance (8.87±1.56 pF; N=57) compared to 3D (7.03±0.59 pF; N=78) and NGN2 (5.26±0.39 pF; N=74) neurons (ANOVA p=0.0008) (**Figure 6a**), suggesting that the cell bodies of dissociated 2D neurons are larger. The membrane resistance of NGN2 (4.69 ±0.43 GΩ; N=74) neurons was higher compared to 2D (2.68±0.26 GΩ; N=57) and 3D (2.94 ±0.16 GΩ; N=78) neurons (ANOVA p<0.0001) (**Figure 6b**), which may indicate fewer ion channels are expressed on the surface of NGN2 neurons. The series resistance of NGN2 (13.27 ± 1.46 MΩ; N=74) neurons was lower than that of 2D (18.41 ± 1.1 MΩ; N=57) and 3D (17.36 ± 0.79 MΩ; N=78) neurons (ANOVA p<0.002) (**Figure 6c**), indicating that a better voltage clamp is achieved in NGN2 neurons.

**Figure 6.**
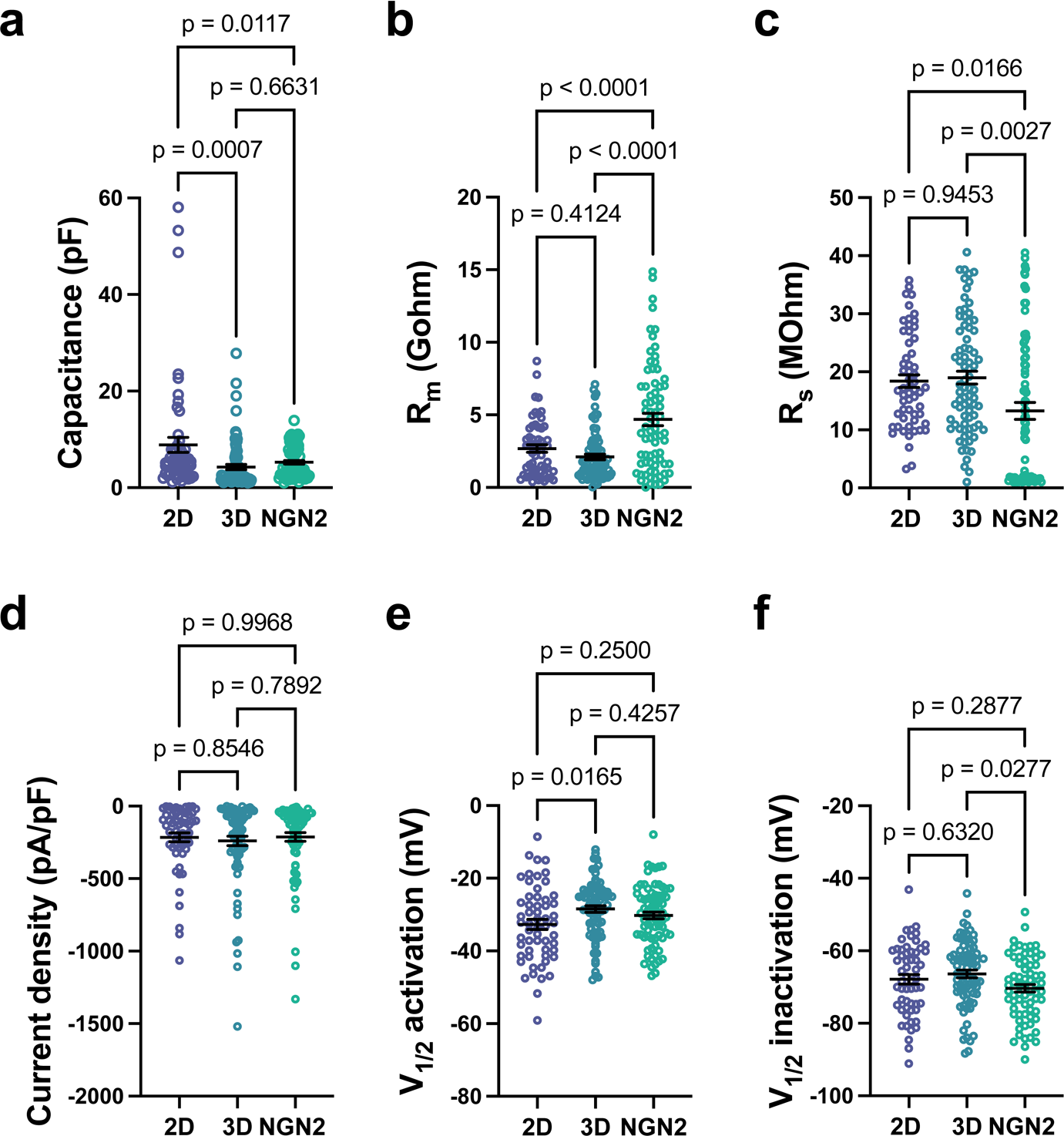
Comparison of membrane and VGSC properties among 2D, 3D, and NGN2 neurons. **(a)** Summary plot comparing capacitance measurements among 2D (8.87±1.56 pF (N=57)), 3D (4.26±0.55 pF (N=78)), and NGN2 (5.26±0.39 pF, (N=74)) neurons (ANOVA p=0.0008). The capacitance of 2D neurons was larger than 3D (p=0.0007) and NGN2 (p=0.0117) neurons. **(b)** Summary plot comparing membrane resistance among 2D (2.10±0.19 GΩ (N=78)), 3D (2.10±0.19 GΩ (N=78)), and NGN2 (4.69±0.43 GΩ (N=74)) neurons (ANOVA p<0.0001). The membrane resistance of NGN2 neurons was larger than 2D (p<0.0001) and 3D (p<0.0001) neurons. **(c)** Summary plot comparing series resistance among 2D (18.41±1.1 MΩ (N=57)), 3D (19.00±1.12 MΩ (N=78)), and NGN2 (13.27±1.46 MΩ (N=74)) neurons (p<0.0001). The series resistance of NGN2 neurons was reduced compared to 2D (p=0.0166) and 3D (p=0.0027) neurons. **(d)** Summary plot showing no difference in the VGSC peak current density among 2D (−215.4±30.4 pA/pF (N=57), 3D (−239.8±32.62 pA/pF (N=78), and NGN2 neurons (−211.90 ±29.96 pA/pF (N=74). **(e)** Summary plot comparing the V_1/2_activation voltage of VGSCs in 2D (V_1/2_= −28.45±0.90mV (N=78)), 3D (V_1/2_=-28.45±0.90mV (N=78)), and NGN2 (V_1/2_=-30.24±0.96mV (N=74)) neurons (ANOVA p=0.0227). The V_1/2_ activation of VGSCs in 2D neurons was more hyperpolarized compared to 3D (p=0.0165) neurons. **(f)** Summary plot comparing the V_1/2_inactivation voltage of VGSCs in 2D (V_1/2_=-67.84±1.30 mV (N=57)), 3D V_1/2_=-66.36±1.06 mV (N=76)), and NGN2 (V_1/2_=-70.34±1.05 mV (N=69)) neurons (ANOVA p=0.0355). The V_1/2_ inactivation of VGSCs in 2D neurons was more hyperpolarized compared to 3D (p=0.0165) neurons.

Comparisons of VGSC properties showed no difference in the average peak current density (ANOVA p=0.78) among 2D neurons (215.4±30.4 pA/pF; N=57), 3D (−239.8±32.62 pA/pF; N=78) neurons, and NGN2 neurons (−211.90±29.96 pA/pF; N=74) (**Figure 6d**). However, the voltage dependence of activation (ANOVA p=0.023) (**Figure 6e**) and inactivation (ANOVA p=0.036) (**Figure 6f**) showed slightly different values among the three models, which is likely due to differences in series resistance, current magnitude ^19^, and maturation ^20^.

### Comparisons between automated and manual patch-clamp

The APC system requires cells to be dissociated, a process which disrupts dendrites and axons, leaving only the cell body intact. This is an important distinction from manual patch-clamp studies, which typically record from intact neurons, and is an important consideration when designing experiments and drawing conclusions from the data generated with this APC system. To begin to determine the effect of neuronal dissociation, we made comparisons of the VGSC activation properties between manual and automated patch-clamp across all three neuronal models. We observed that the VGSC V_1/2_ activation voltage was similar across manual and automated patch-clamp within each neuronal model (**Figure 7**). V_1/2_ inactivation was not compared, as it can be influenced by current amplitude ^19^, which is typically larger when using manual patch-clamp due to experimenter bias related to selecting optimal neurons to record. These results suggest that for measuring VGSC activation, the high-throughput and unbiased nature of APC is advantageous compared to manual patch-clamp.

**Figure 7.**
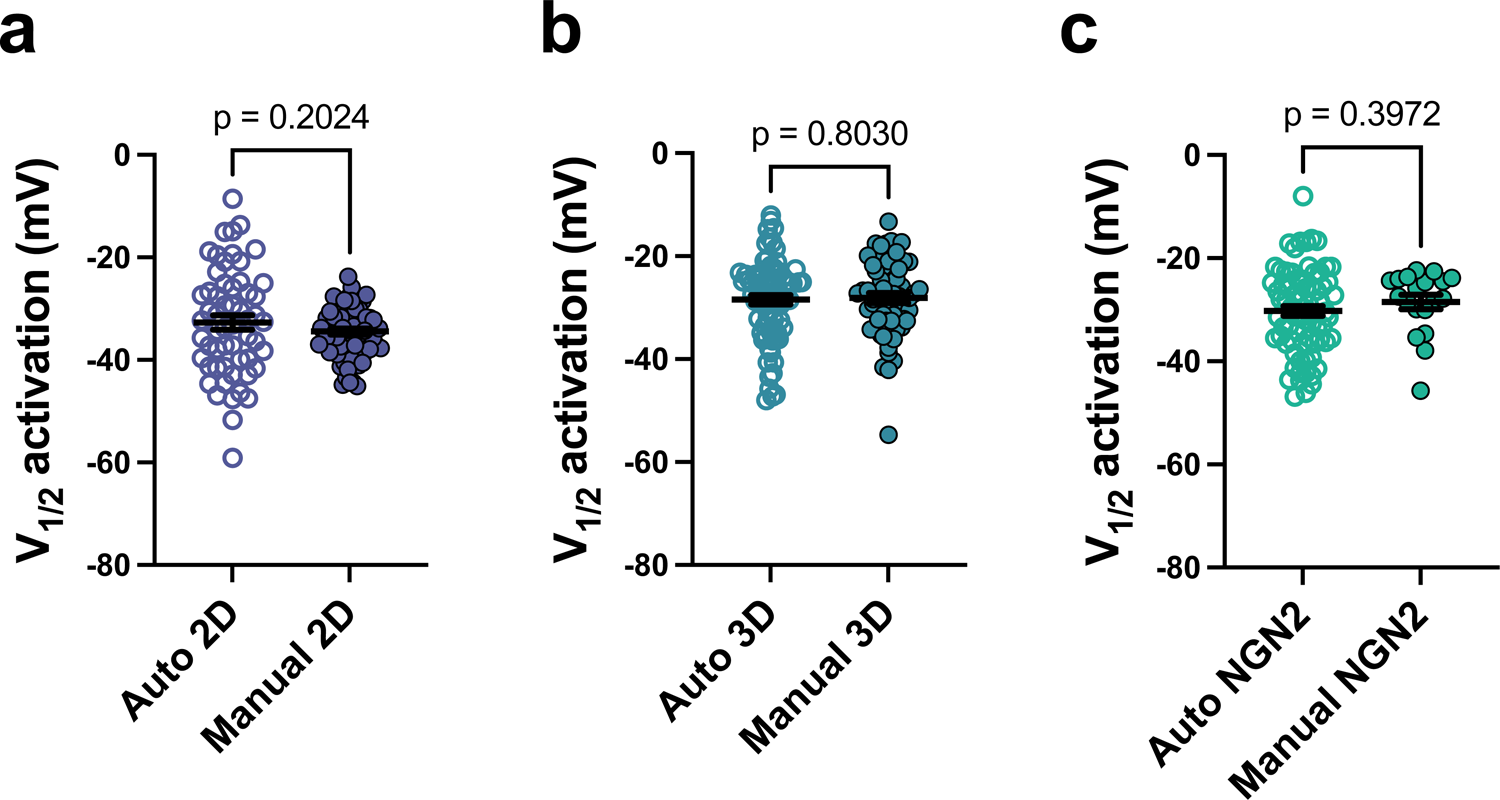
Automated patch-clamp recording produces similar results compared to manual patch-clamp. **(a)** Summary data comparing the VGSC V_1/2_ activation of 2D neurons obtained between automated and manual patch-clamp (Auto 2D V_1/2_=-32.7±1.4 mV (N=57), Manual 2D V_1/2_=34.5±0.6 mV (N=74); unpaired t-test p=0.2024). **(b)** Summary data comparing the VGSC V_1/2_ activation of 3D neurons obtained between automated and manual patch-clamp (Auto 3D V_1/2_=-28.45±0.90 mV (N=78), Manual 3D V_1/2_=-28.12±0.96 mV (N=60); p=0.8030). **(c)** Summary data comparing the VGSC V_1/2_ activation of NGN2 neurons obtained between automated and manual patch-clamp (Auto NGN2 V_1/2_=-30.24±0.96 mV (N=74), Manual 3D V_1/2_=-28.51±1.40 mV (N=19); p=0.3972).

### Pharmacology of VGSCs with automated patch-clamp

An important aspect of drug discovery is the ability to perform compound screening on physiologically relevant cells or tissue. Pharmacology experiments using manual patch-clamp are labor intensive and low throughput. Typically, a single compound at a single concentration is tested on a single recorded neuron, and compounds are not easily washed out, which necessitates using a fresh experimental prep after every drug application. Moreover, acquiring useful data is challenging because it depends on maintaining stable recordings. For APC systems, pharmacological approaches have only been possible in heterologous cells expressing recombinant ion channels, which limits its translational potential because these cell systems lack the accessory proteins and biochemical pathways that are known to regulate ion channel function^21^. To demonstrate the feasibility of pharmacological screening of native ion channels in human neurons we tested the inhibition of VGSC currents with XPC6444, a highly potent inhibitor of Nav1.6 (IC50=41nM) and Nav1.2 (IC50=125nM) ^22^. We recorded baseline VGSC currents from dissociated organoids derived from two hiPSC lines and then delivered XPC6444 (250nM) using the liquid handling robot on the APC system. XPC6444 blocked 27% of the max current density across both lines (baseline −119.4±2.06 pA/pF, XPC6444 −86.9±13.8 pA/pF; paired t-test p=0.0006; N=45) (**Figure 8a,b**). We also observed that application of XPC6444 produced a hyperpolarized shift in the V_1/2_ activation of VGSC currents (baseline −28.4±1.1 mV, XPC6444 - 30.6±1.1 mV; paired t-test p=0.0029; N=45) (**Figure 8c, d).** Together, these results confirm the presence of Nav1.6 and/or Nav1.2 isoforms in hiPSC-derived neurons dissociated from cortical organoids and demonstrate the suitability of automated patch-clamp for performing pharmacology and compound screens on native ion channels expressed in human neurons.

**Figure 8.**
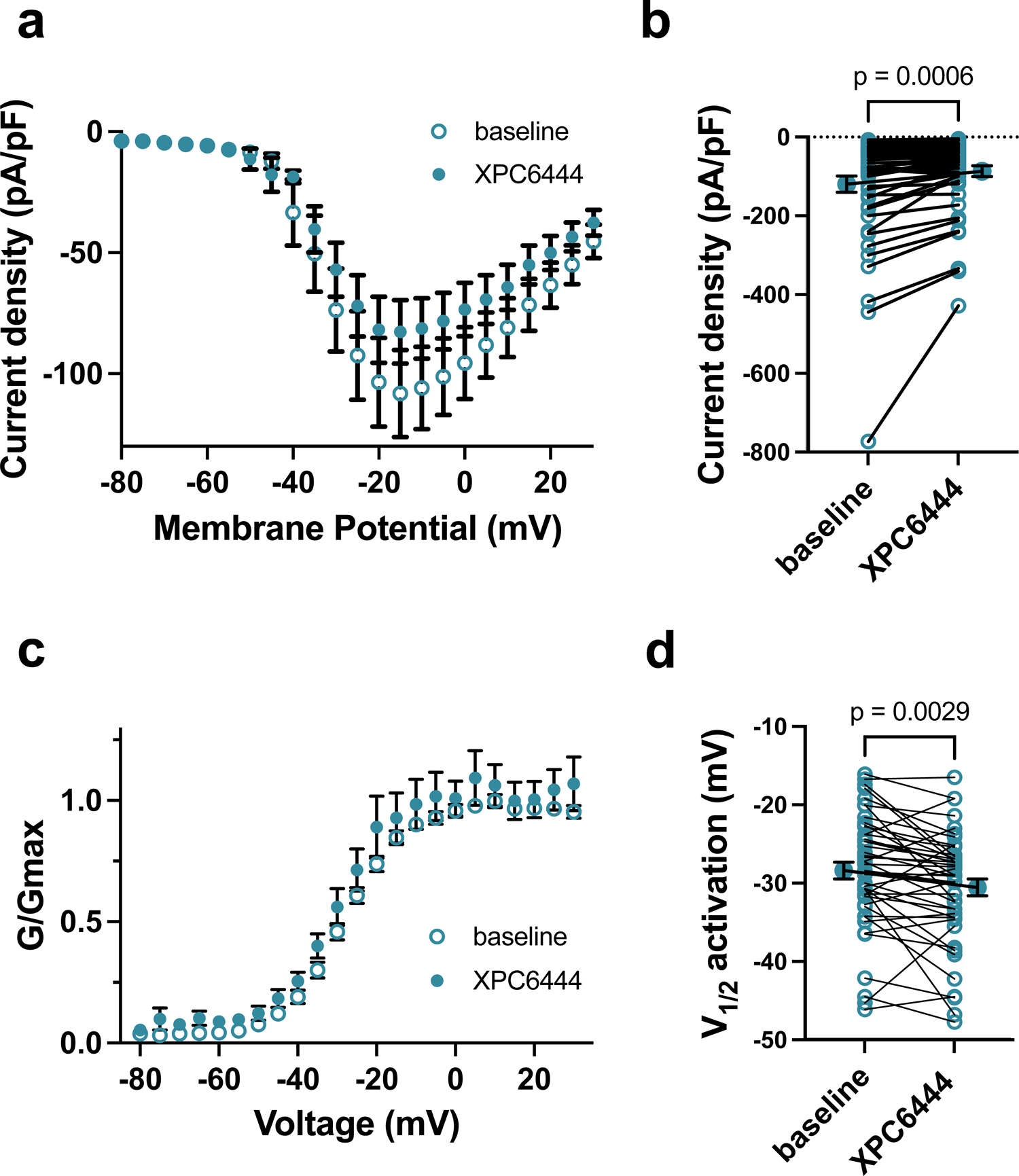
VGSC pharmacology in 3D neurons using the automated patch-clamp system. **(a)** Current-voltage plot of VGSC currents before and after application of XPC6444 (N = 48 recordings). **(b)** Summary data showing the application of XPC6444 blocks a portion of the VGSC current (baseline −119.4±2.06 pA/pF, XPC6444 −86.9±13.8 pA/pF; paired t-test p=0.0006 (N=45). **(c)** Voltage-dependence of steady-state activation before and after application of XPC6444. **(d)** Summary data showing application of XPC6444 leads to a hyperpolarized shift in the V_1/2_activation of VGSCs in 3D neurons (baseline −28.4±1.1 mV, XPC6444 −30.6±1.1 mV; paired t-test p=0.0029 (N=45)).

### FACS prior to automated patch-clamp

To further enhance versatility and to assess the relationship between the success rate and cell-type complexity in hiPSC-derived models using APC system, we next tested the addition of a FACS step into our APC workflow. We reasoned that the reduced recording success rate from 3D neurons could be attributed to the cell-type complexity inherent to 3D organoids. We virally infected organoids at day 138 with a Syn1-GFP reporter for 48h and after 10 days we dissociated them and performed FACS to enrich for Syn1+ neurons prior to APC recording. For this comparison, we dissociated organoids from the same hiPSC line without FACS, equalized cell density between no FACS and FACS (17K/mL), and performed side-by-side APC recordings within the same 384-well plate. We recorded VGSC currents from FACS 3D neurons and obtained a significantly higher success rate compared to non-FACS 3D neurons (**Figure 9**). These results suggest that the lower recording success rate observed in uninfected dissociated 3D neurons (**Figure 9b**) can be partly attributed to the cell-type complexity of organoids, which can be resolved by adding a FACS step to enrich for neurons prior to APC recording. In addition, this experiment demonstrates the feasibility of recording from genetically manipulated cellular populations using cell-type specific reporters and/or CRISPR/Cas9 technologies, which greatly enhances the experimental repertoire of this approach.

**Figure 9.**
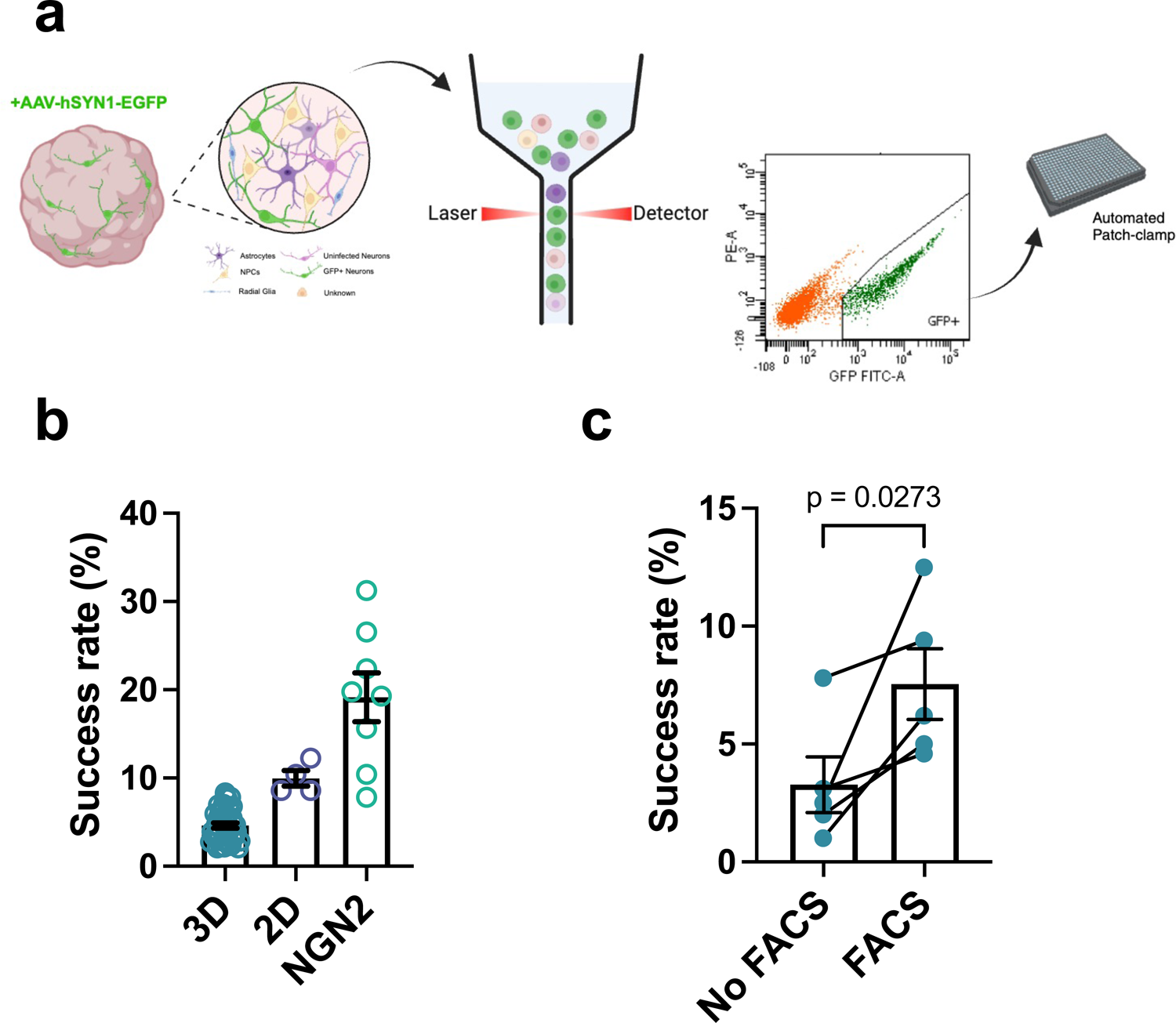
FACS improves the APC recording success rate. **(a)** Workflow diagram. Lenti-Syn1-GFP infected 3D organoids are dissociated and FACS. GFP+ cells are then dispensed onto the APC platform. **(b)** Summary data showing APC recording success rates (% of wells) for 3D (4.62±0.29, N=41), 2D (9.96±0.87, N=4), and NGN2 (19.14±2.77, N=8). **(c)** Summary data showing the recording success rate for 3D organoids is increased after FACS (No FACS 3.28±1.18, FACS 7.54±1.5; paired t-test p=0.027 (N=5)).

## DISCUSSION

Patch-clamp electrophysiology was originally developed by Sakmann and Neher, and revolutionized the fields of physiology and neuroscience by enabling the resolution of currents from single ion channels^23,24^. Further advances of this technique have led to our broad understanding of excitable cells, from the biophysical properties of single ion channels in a patch of membrane to whole-cell measures of action potentials and synaptic transmission. The primary limitations of this technique are throughput and accessibility, as it requires specialized equipment and expertise to place an electrode onto a single cell. The development of automated patch-clamp has overcome some of these limitations, but until recently, it has primarily been applied to studying heterologous cells expressing recombinant ion channels^4^. Adaptation of automated patch-clamp for excitable cells has begun with several groups demonstrating its use to study cardiomyocytes, hiPSC-derived dopaminergic neurons, and primary cortical and dorsal root ganglion neurons ^25–28^. Our results expand on these prior studies by providing methods and technological advances for automated recording of three different, widely used, hiPSC-derived models of the cortex (2D, 3D, NGN2). We anticipate that the combination of this APC method along with hiPSC-derived models will help to overcome the bottlenecks in throughput related to manual patch-clamp and human genetic variation.

Across the three models we tested, our success rate obtaining recordings across the 384 wells varied between 5% (3D), 10% (2D) and 19% (NGN2) depending on the model type. These percentages are low compared to APC recording from heterologous cells expressing recombinant ion channels (approx. 80%)^29^. However, even a 5% success rate equates to obtaining electrophysiology data from 19 cells in 30 minutes, which would take a highly trained electrophysiologist at least two days to accomplish via manual patch-clamp. We attribute this large range of success rates primarily to the cellular and structural complexity of the hiPSC model being studied. Our lowest success rate was obtained from dissociated 3D organoids which are composed of a variety of non-neuronal cell types such as radial glial cells, intermediate neuroprogenitor cells, and astrocytes ^30^, all of which are dispensed across the 384 wells of the APC system. Our 2D model, which contains human cortical neurons and rat astrocytes in approximately a 1:5 ratio, produced an intermediate success rate, and our NGN2 model, which is a relatively pure population of neurons, obtained the highest success rate. In addition, an important prerequisite of patch-clamp recordings is healthy cellular membranes, and we speculate that dissociation of tissue (3D) compared to monolayers of cells (2D, NGN2) results in more cellular debris and reduced membrane health that further contributes to our reported successful recording ranges. The effect of these two variables (cell types and health/debris) on success rate was demonstrated in our FACS experiment (**Figure 9**), whereby the enrichment of neurons and removal of cellular debris significantly improved the recording efficiency from 3D organoids.

An important consideration when deciding between manual patch clamp and APC experiments relates to the question being asked. Cellular dissociation is a prerequisite for APC experiments which leads to the disruption of neuronal connectivity and cleavage of dendrites and axons, leaving only the neuronal cell body intact. Therefore, APC recording is not suitable for assaying network connectivity, synaptic transmission, and the biophysical properties of channels, transporters and receptors that are localized to axons and dendrites. However, the membranes of the neuronal soma and somatodendritic regions are highly enriched in a variety of voltage - gated ion channels which can be studied using APC systems. Given the significant morphological differences between intact neurons and dissociated neurons, which will affect intrinsic membrane properties and current amplitudes, we only compared VGSC steady-state activation between manual and APC (**Figure 7**). Across all three hiPSC models, we observed a similar distribution of V_1/2_ activation voltages. At the biophysical level, these results are not surprising because the cable properties of dendrites and axons are known to limit the amount of the neuron that is effectively voltage-clamped ^31^ and VGSCs are highly localized to the soma and somatodendritic regions of neurons ^32^. These consistencies suggest that the technical differences associated with APC recording do not appear to significantly degrade measurement of VGSC activation. However, this may not be the case for other voltage-gated channels, such as potassium and calcium channels, which may have more varied distributions across neuronal compartments ^32^.

Human neurons derived from hiPSCs are relatively immature, based on their physiological characteristics and transcriptional profiles. Neurons from the 3D organoid protocol we employed were shown to be comparable to fetal brain between 10 and 24 postconceptional weeks (PCW), based on comparisons of methylation and transcription between 3D hCOs and fetal brain ^16^. Cortical neurons derived from our 2D protocol were shown to be transcriptionally equivalent to late third-trimester fetal brain ^33^. The transcriptome of 28-day NGN2 neurons indicated they are approximately between 8 and 24 PCW ^34^, however, this study added dual SMAD inhibition to their protocol, which is a modification from the protocol we used. With consideration of the known maturational differences among these three protocols, we made comparisons of their electrophysiological properties (**Figure 6**). We observed that 2D neurons had larger capacitance compared to 3D and NGN2, and NGN2 neurons displayed larger membrane resistance compared to 2D and 3D neurons. These differences in intrinsic membrane properties corresponded well with the differences in transcriptional maturation.

A major advantage of the high-throughput capabilities of APC recording is its applicability to pharmacology and drug screening^35^. For instance, every new drug requires electrophysiological testing against hERG channels expressed in heterologous cells for cardiac safety ^36^, and recently a 1920 compound screen for allosteric modulators of NMDA receptors was reported ^37^. One potential way to improve translation in drug development is to perform drug screens and pharmacology on native ion channels and receptors in human cell types. Native channels are associated with large signaling complexes that regulate channel kinetics, trafficking, and anchoring ^38–40^, all of which can affect how compounds ultimately modulate channel function in vivo. Here, we demonstrate the suitability of APC for pharmacology studies on native VGSC in human neurons (**Figure 8**). We are the first to demonstrate that VGSC currents in neurons from 3D organoids are sensitive to XPC-6444, a highly potent, isoform-selective inhibitor of Nav1.6 and Nav1.2, which indicates that one or both of these VGSC contributes approximately 27% of the VGSC currents in such 3D neurons. These results highlight the potential of combining hiPSC-derived neuronal models and APC for high-throughput drug screening and pharmacology of native channels in human neurons.

To improve our ability to elucidate the contribution of specific ion channels and their associated molecular complexes on cellular excitability, it is important to develop methods to manipulate specific proteins associated with channel complexes in their native environment. This will enable us to identify specific accessory proteins that interact with channels and how they modulate channel function, thus allowing for the identification of novel drug targets. An additional challenge associated with ion channel drug development is the significant sequence homology among human ion channel isoforms, which makes it difficult to develop compounds that are selective ^41^. The specificity of genetic manipulation is a viable approach to elucidate the contribution of accessory proteins and overcome issues related to ion channel similarity. To enable genetic manipulation approaches, we added a FACS step into our APC workflow (**Figure 9**). We found that enrichment of Syn1+ neurons via FACS improved recording efficiency and confirmed the ability to record from genetically manipulated neurons. This approach can easily be adapted to CRISPR/Cas9 technology to target specific genes, which will enhance functional studies investigating the role of ion channels and accessory proteins on neuronal excitability. In addition, this approach can be combined with cell type-specific reporters enabling the enrichment of specific subtypes of excitable cells.

The advent of hiPSC models opens up an exciting new research avenue that is rapidly improving our ability to model human genetic variation and disease, which should improve the fidelity of translation from the bench to the bedside. To realize the immense translational potential of hiPSC models, innovative high-throughput methods are needed to understand and leverage the immense complexity of human genetics and disease. By combining three commonly used hiPSC models of brain development with APC recording, we have established an innovative approach to assay native ion channels in human neurons. The high-throughput capabilities of APC allow for unbiased and simultaneous comparisons across many human genomes, compounds, and/or genetic manipulations, which we anticipate will accelerate our understanding of human disease and the development of therapeutics.

## AUTHOR CONTRIBUTION

**F. Farinelli:** Methodology, Investigation, Validation, Formal analysis, Data curation, Writing - Original Draft, Writing - Review & Editing**. I. Ostlund:** Methodology, Investigation, Writing-Original Draft. **S.R. Sripathy:** Methodology, Investigation, Visualization, Writing - Original Draft, Writing - Review & Editing. **D. Das:** Methodology, Investigation, Writing - Original Draft, Writing - Review & Editing. **G. Shim:** Methodology, Investigation, Writing - Review & Editing. **S. Myung:** Methodology, Investigation. **R.E. Straub:** Writing - Review & Editing, Funding acquisition. **B.J. Maher:** Conceptualization, Methodology, Formal Analysis, Resources, Data Curation, Visualization, Writing - Original Draft, Writing - Review & Editing, Supervision, Project administration, Funding acquisition.

## ACKNOWLEDGEMENTS

We are grateful for the vision and generosity of the Lieber and Maltz families, who made this work possible. This project was supported by the Lieber Institute for Brain Development, the National Institute of Mental Health (Grant No. R01MH134828 (to B.J.M, R.E.S), Grant No. R01MH110487 (to BJM), Grant No. R21MH132035 (to BJM). The collection of fibroblasts, from which the hiPSCs were derived, was supported by direct funding from the Intramural Research Program (IRP) of the NIMH to the Clinical Brain Disorders Branch (D.R.W., PI, Protocol 95-M-0150, NCT00001486, Annual Report no. ZIA MH002942053) with supplemental support from the Clinical and Translational Neuroscience Branch (Karen Berman, PI). We thank all of the participants in the IRP study and their families. The content is solely the responsibility of the authors and does not necessarily represent the official views of the National Institutes of Health.

